# IL-4 Licenses B Cell Activation Through Cholesterol Synthesis

**DOI:** 10.1101/2024.05.13.593964

**Authors:** Holly R. Steach, Autumn G. York, Jonathan Mattingly, Mathias H. Skadow, Shuting Chen, Jun Zhao, Kevin J. Williams, Quan Zhou, Weu-Yuan Hsieh, J. Richard Brewer, Rihao Qu, Justin A. Shyer, Christian Harman, Esen Sefik, Walter K. Mowell, Aimee Wu, Meijie Aletta Li, Will Bailis, Can Cui, Yuval Kluger, Steven J. Bensinger, Joe Craft, Richard A. Flavell

## Abstract

Lymphocyte activation involves a transition from quiescence and associated catabolic metabolism to a metabolic state with noted similarities to cancer cells such as heavy reliance on aerobic glycolysis for energy demands and increased nutrient requirements for biomass accumulation and cell division^1–3^. Following antigen receptor ligation, lymphocytes require spatiotemporally distinct “second signals”. These include costimulatory receptor or cytokine signaling, which engage discrete programs that often involve remodeling of organelles and increased nutrient uptake or synthesis to meet changing biochemical demands^4–6^. One such signaling molecule, IL-4, is a highly pleiotropic cytokine that was first identified as a B cell co-mitogen over 30 years ago^7^. However, how IL-4 signaling mechanistically supports B cell proliferation is incompletely understood. Here, using single cell RNA sequencing we find that the cholesterol biosynthetic program is transcriptionally upregulated following IL-4 signaling during the early B cell response to influenza virus infection, and is required for B cell activation *in vivo*. By limiting lipid availability *in vitro*, we determine cholesterol to be essential for B cells to expand their endoplasmic reticulum, progress through cell cycle, and proliferate. In sum, we demonstrate that the well-known ability of IL-4 to act as a B cell growth factor is through a previously unknown rewiring of specific lipid anabolic programs, relieving sensitivity of cells to environmental nutrient availability.

## IL-4-Dependent Phase of Early Antiviral B Cell Activation Requires Cholesterol Synthesis

To investigate whether Il-4 engages specific metabolic programs to support B cell proliferation during physiologically relevant activation, we focused on the early B cell response to influenza virus infection, which is dependent on IL-4 production by NK-T cells at interfollicular sites in the mediastinal lymph node^8^. We performed single cell RNA sequencing (scRNAseq) on total CD19+ B cells sorted from mediastinal lymph nodes 4 days post-infection with PR8 influenza. To determine how specific transcriptional programs mapped to the developmental trajectory of B cell activation, we employed semi-supervised trajectory inference based on expression of the quiescence gene Btg2 and activation gene Myc using Monocle^9–11^. Pseudotime trajectory reconstruction enabled visualization of the gradual progression from naïve to activated states (Fig.1a-c). As expected, progression across pseudotime was marked by loss of quiescence gene expression (Btg2, Tsc22d3) and upregulation of transcripts associated with activation and cell cycle, indicating proliferation (Cd86, Ccnd2, Myc) (Fig.1b,c). By analyzing the expression of specific genes of interest across pseudotime, we were able to resolve where select transcriptional programs fit into the coordinates of B cell activation. We observed a distinct IL-4 signature in a subset of activated B cells marked by expression of Il4i1 confirming recent findings that B cells require IL-4 during early responses to influenza (Fig.1a,b)^8^. We additionally found a strong association between this IL-4 signature and transcription of genes involved in cholesterol synthesis (eg. Sqle, Hmgcr, Mvk, Dhcr7; Fig.1b,c).

Cholesterol biosynthesis is regulated by a master transcription factor sterol-regulatory element binding protein 2 (SREBP2)^12^. SREBP2, encoded by the gene Srebf2, is held in the ER by chaperones SCAP and INSIG, both of which are directly regulated by sterols present in the ER membrane. When sterol levels are low, INSIG dissociates from SCAP/SREBP2 permitting their translocation to the Golgi apparatus where SREBP2 is cleaved and activated by site 1 and 2 proteases (S1P and S2P, respectively; extended data Fig.1a). Active SREBP2 translocates to the nucleus to mediate transcription of genes involved in the cholesterol biosynthetic pathway such as Hmgcr, Sqle, and Dhcr24. This program is subject to feedback inhibition whereby synthesized cholesterol accumulates in the ER and associates with SCAP and INSIG to prevent Golgi trafficking and activation of SREBP2. Additionally, the intermediate metabolite desmosterol ligates nuclear hormone Liver X Receptors (LXRs), which drive transcription of genes involved in efflux of cholesterol out of cells such as Abca1, Abcg1, and Apoe13. Coordinate regulation of these positive and negative transcriptional programs together mediates metabolic homeostasis. Cytokine signaling and other immune cell activation pathways have been shown to influence de novo lipid synthesis and export programs in T cell and macrophage^13–16^; however, the signals that regulate how B cells balance opposing arms of cholesterol homeostasis in response to different stimuli are largely unclear. Although SREBP2 activation often leads to a coordinate upregulation of LXR target genes by engaging a counter-regulatory program, we surprisingly observed a decrease in LXR target genes (Abca1, Apoe; Fig.1b,c) co-occurring with the observed upregulation of SREBP2 target genes (Hmgcs1, Hmgcr, Mvk, Sqle, Dhcr24) coincident with an IL-4 signature. We performed qPCR at various timepoints following influenza virus infection to determine whether global gene expression patterns matched the kinetics across activation defined pseudotime (Fig. 1.d). We observed a marked upregulation in SREBP2 target genes (Hmgcs1, Hmgcr, Mvk, Sqle, Dhcr24) and coordinate downregulation of LXR target (Abca1, Abcg1) genes beginning at the peak of IL-4 production by NK-T cells (Fig.1d). Together, these data suggest that IL-4 signaling may support early B cell proliferation by tuning the balanced regulation of cholesterol homeostasis to promote synthesis via regulation of SREBP2 and accumulation by limiting cellular efflux.

**Figure 1.**
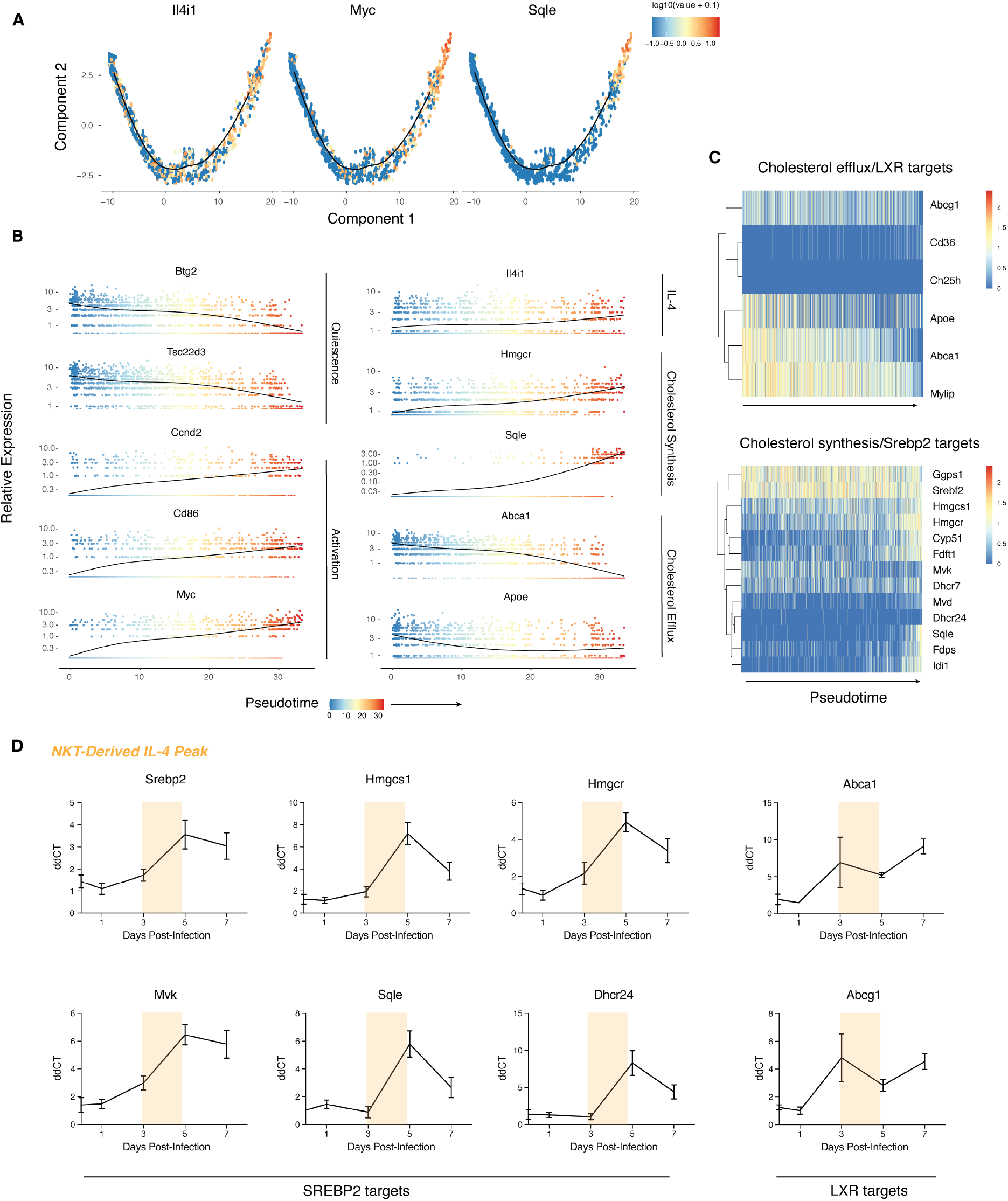
IL-4-dependent early antiviral B cell activation requires cholesterol synthesis. **a**, Activation-defined pseudotime trajectory of scRNAseq profiles of murine B cells on day 4 of PR8 influenza virus infection colored by relative expression of Il-4 signature, activation, and cholesterol synthesis genes **b**, Kinetics plot showing relative expression of activation, quiescence, IL-4 signature, cholesterol synthesis, and cholesterol efflux genes across activation pseudotime colored by pseudotime value **c**, Heatmap showing Log-normalized expression of genes involved in cholesterol efflux and cholesterol synthesis across activation pseudotime **d**, Gene expression of isolated B cells analyzed at various timepoints through early infection with influenza virus. Colored section indicates the peak of IL-4 production by NK-T cells as previously defined^8^

These data show a critical role for cholesterol synthesis during early B cell activation, suggesting a direct cellular mechanism through which statins may impact humoral immunity. Sequential phases of B cells activation in secondary lymphoid tissues are highly spatially compartmentalized, leading to dense areas of actively dividing cells^17^. This likely results in local competition for limiting nutrients. We therefore hypothesized that B cell proliferation in response to mitogenic stimuli such as antigen recognition requires cholesterol, and that in the absence of environmental lipids IL-4 may compensate and enable proliferation via upregulation of endogenous cholesterol synthesis.

## IL-4 Circumvents A Metabolic Requirement for Exogenous Cholesterol via Upregulation of Endogenous Synthesis via SREBP2 and Stat6

To determine the necessity for IL-4 mediated cholesterol synthesis in early B cell expansion in conditions where extracellular nutrients are limiting, we developed an *in vitro* system whereby CellTrace™ Violet-labeled B cells were stimulated with anti-IgM and anti-CD40 for 96 hours in a minimal media formulated to contain non-nutrient components of serum including insulin, transferrin, selenium, ethanolamine, and albumin but, importantly, no lipids. Relative to controls stimulated in 10% Fetal Bovine Serum, lipid deprivation caused profound defects in proliferation that was strikingly rescued by treatment with IL-4 or, in a dose-dependent manner by the addition of M*β*CD-solubilized cholesterol (Fig.2a-d). Importantly, supplementation of either BSA-conjugated palmitate (16:0; a saturated fatty acid) or oleic acid (18:1; a monounsaturated fatty acid) were unable to produce the same rescue (Fig.2e,f), suggesting that cholesterol is uniquely required for B cell activation and proliferation.

**Figure 2.**
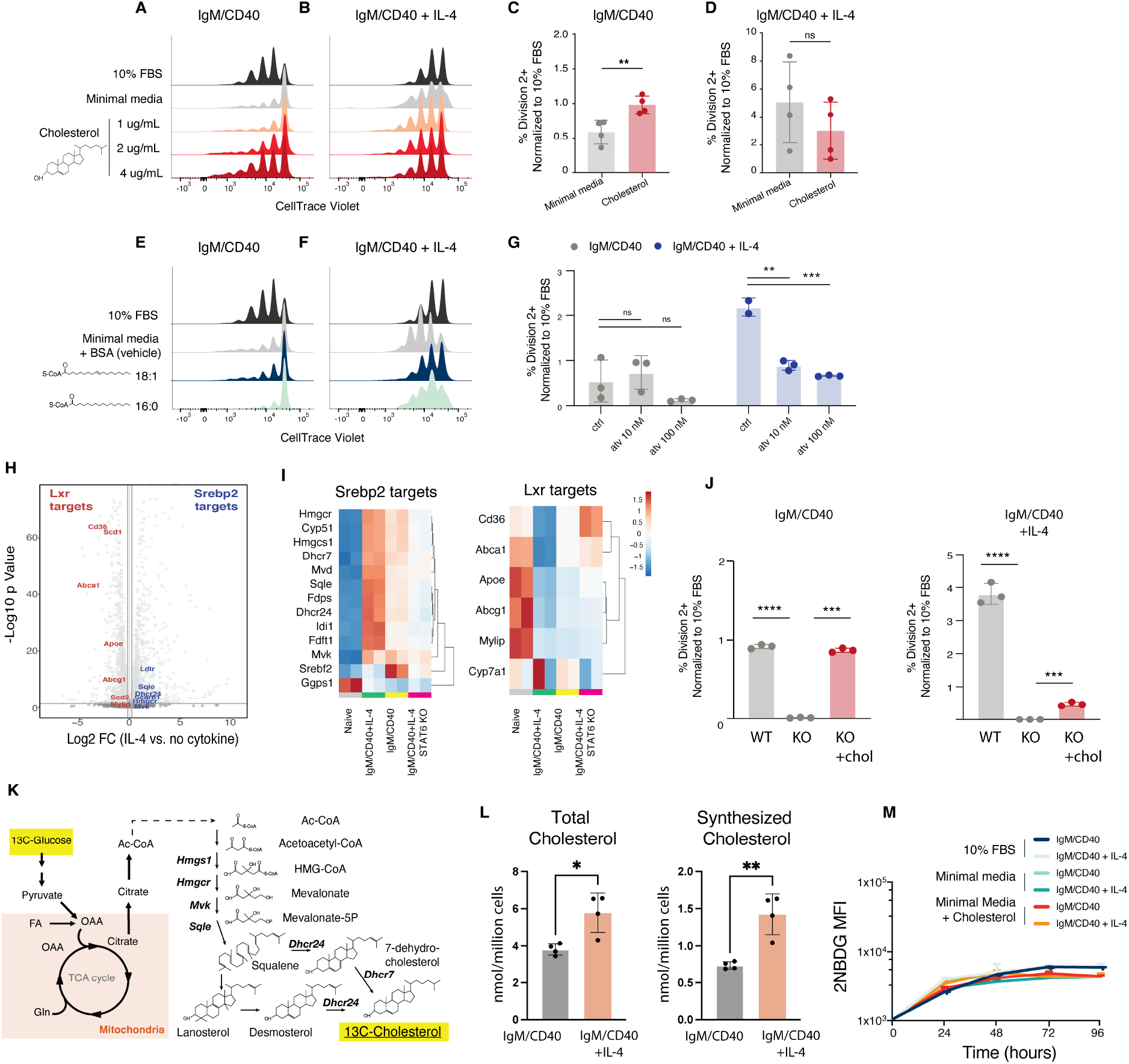
IL-4 circumvents a metabolic requirement for exogenous cholesterol via upregulation of endogenous synthesis. **a**,**b**, Representative plots and **c**,**d**, Summary data of B cell proliferation following 96H stimulation minimal media supplemented M*β*CD-solubilized cholesterol, or 10% FBS and minimal media controls. Mean and S.D. from 5 independent experiments are represented on summarized plots; each dot represents an average of 3 replicates within a single experiment. **P = 0.01, unpaired two-tailed Student’s t-test comparing Lipid Free and Cholesterol treated B cells **e**,**f** Proliferation of B cells stimulated for 96H in minimal media supplemented BSA-conjugated fatty acids, or 10% FBS and BSA vehicle as controls **g**, Summary of B cell proliferation following 96H stimulation with anti-IgM, anti-CD40, +/-IL-4 in Minimal Media in the presence of DMSO or Atorvastatin. Mean and S.D. from 3 biological replicates are represented on summarized plots. **P = 0.01, unpaired two-tailed Student’s t-test. All experiments were performed at least twice. **h**, Volcano plot summarizing differential expression of genes measured by RNA sequencing in B cells stimulated for 72H in varying media conditions with anti-IgM/CD40 +/-IL-4 **i**, Heat map summarizing gene expression in wild type or STAT6 knockout B cells stimulated for 72H in varying media conditions with anti-IgM/CD40 +/-IL-4 **j**, Summary data of Mb1-Cre SREBP2 fl/fl or Mb1-Cre control B cells stimulated with or without IL-4 in minimal media supplemented M*β*CD-solubilized cholesterol. Mean and S.D. from 3 biological replicates are represented on summarized plots. **P = 0.01, unpaired two-tailed Student’s t-test. All experiments were performed at least twice. **k**, Graphical illustration of glucose-derived 13C incorporation into actively synthesized cholesterol **l**, Absolute amount of total (left) and synthesized (right) cholesterol as measured by Isotopic Spectral Analysis (ISA) following 72H stimulation as indicated. Mean and S.D. representing 4 technical replicates on summarized plots. *P = 0.05, **P = 0.01, ***P < 0.001, ****P < 0.0001, unpaired two-tailed Student’s t-test used to determine significance. Data are representative of two independent experiments. **m**, Glucose uptake measured by 2-NBDG incorporation at 24-hour time intervals

We next assessed the effects of statins, a widely prescribed class of cholesterol-lowering drugs that primarily target the rate-limiting enzyme HMG-CoA reductase (HMGCR), the most abundantly prescribed of which is Atorvastatin (Lipitor®). Various studies have reported mixed results on statins impacting antiviral immunity following influenza vaccination in humans; yet, how this occurs mechanistically is unclear^18–20^. The capacity of IL-4 to rescue proliferation of lipid deprived cells was blocked by Atorvastatin, implying that IL-4 renders cells resistant to external lipid deprivation through upregulation of endogenous cholesterol synthesis.

To verify the transcriptional targets of IL-4 signaling, we performed bulk RNA sequencing on IL-4 stimulated B cells in 10% FBS. These experiments revealed a marked IL-4 dependent upregulation of SREBP2 target genes involved in cholesterol synthesis and downregulation of LXR target genes involved in cholesterol efflux, recapitulating the gene expression pattern observed *in vivo* (Fig.2h,i, Extended Data Fig.1a), which was confirmed by qPCR in both C57BL/6J and C57BL/6N backgrounds (Extended Data Fig.1b,c). We hypothesized that the observed increase in cholesterol synthesis genes maybe require STAT6, a transcription factor that is downstream of IL-4 receptor activation. Indeed, RNAseq analysis of STAT6 knockout cells indicated that STAT6 was required for IL-4 mediated regulation of SREBP2 and LXR target genes (Fig.2i, extended data Fig.1a).

In order to confirm that IL-4 and its effector STAT6 were indeed acting through the canonical master regulator SREBP2, we generated Mb1-Cre Srebf2 flox/flox conditional B cell knockout mice. We observed that, in comparison to Mb1-Cre controls, B cells deficient in SREBP2 were highly insensitive to IL-4 and were unable to proliferate in minimal media; treatment with exogenous cholesterol was able to partially rescue these proliferative defects (Fig.2j).

To verify changes in cholesterol metabolism induced by transcriptional regulation downstream of IL-4 signaling, we employed gas chromatography-mass spectrometry (GC-MS) paired with 13C-stable isotope enrichment studies to track glucose-derived Acetyl-CoA^14–16^ to measure synthesized cholesterol in B cells. In the presence of IL-4, we observed a significant increase in both total and actively synthesized cholesterol (Fig.2k,l). Changes in labelled cholesterol were not due to differences in glucose uptake as measured by uptake of 2NBDG (Fig.2m), demonstrating that IL-4 regulates changes in glucose-derived carbon allocation to support cholesterol biosynthesis. Prior studies have shown that IL-4 increases glucose uptake, however these studies employed naïve B cells treated with cytokine alone, not activated cells receiving IL-4 as an additional signal^21^. Together, these data demonstrate that Il-4 acts as a B cell co-mitogen through upregulation of endogenous cholesterol synthesis via the transcription factors Stat6 and SREBP2.

## IL-4 or Extracellular Cholesterol is Dispensable for Proximal Signaling Downstream of BCR/CD40 but Necessary for ER Expansion and Cell Cycle Entrance

To understand mechanistically how cholesterol and IL-4 regulate B cell proliferation, we assessed their cell cycle status by co-staining with Ki-67 and 7AAD allowing us to partition cells into G0, G1, G2/S, and M phases (Fig.3a,b representative gating Fig.3c). In the absence of exogenous cholesterol or IL-4, we observed a robust accumulation of cells in G0, indicating that these cells were unable to enter cell cycle. Notably, cell cycle progression was rescued to a similar extent by addition of IL-4 or cholesterol, demonstrating a critical IL-4/cholesterol axis in licensing exit of quiescence and activation of B cells.

**Figure 3.**
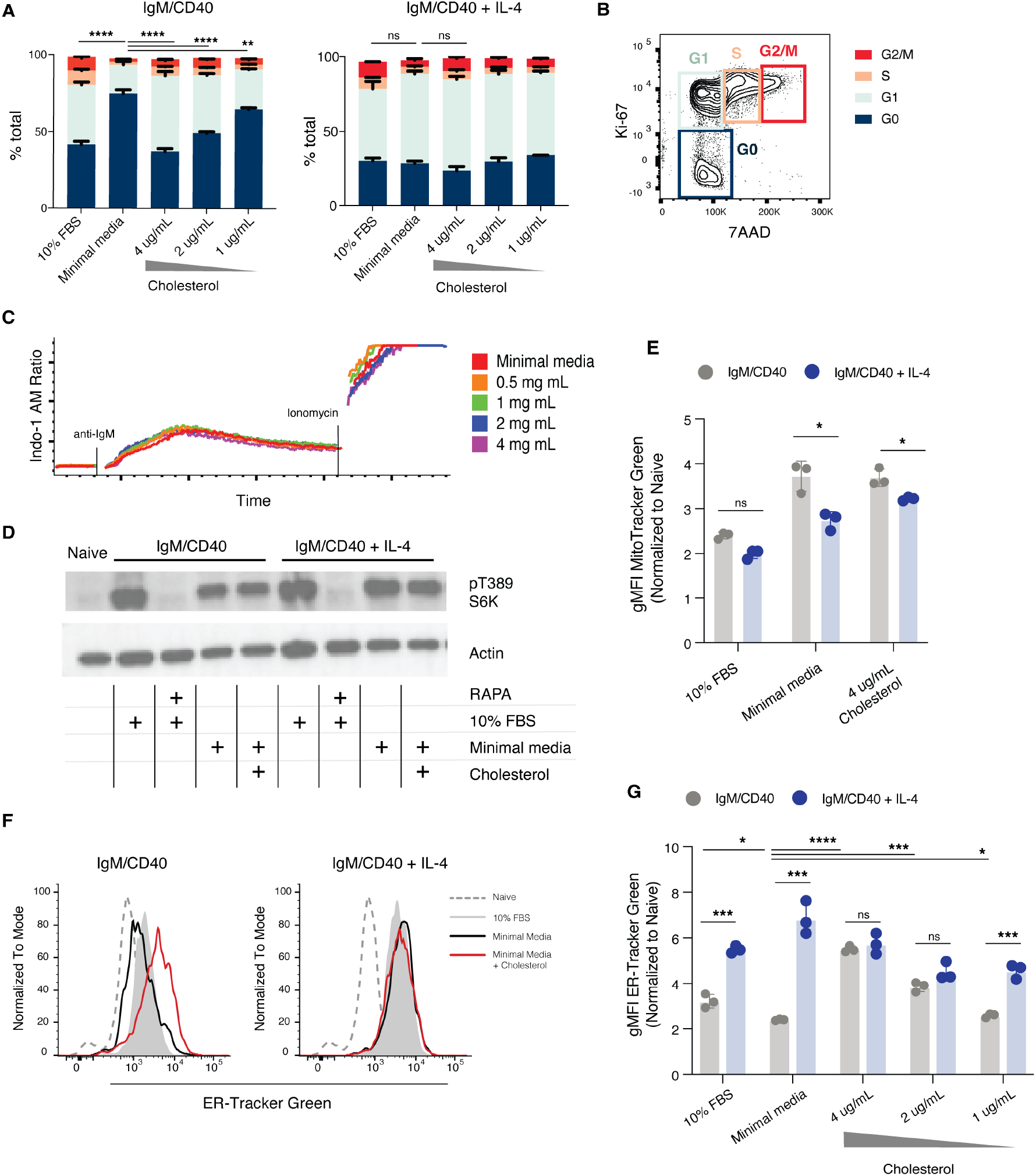
IL-4 or extracellular cholesterol is dispensable for proximal signaling downstream of BCR/CD40 but necessary for ER expansion and cell cycle entrance. **a**, Summary data and **b**, representative gating of cell cycle status following 72H stimulation. Mean and S.D. representing 3 biological replicates on summarized plots. *P = 0.05, **P = 0.01, ***P < 0.001, ****P < 0.0001, unpaired two-tailed Student’s t-test comparing percentage of cells in G0 between Minimal Media and other conditions. All experiments were performed at least twice. **c**, Calcium flux measured in Indo-1 AM-loaded B cells treated with 10 ug/mL anti-IgM Fab2 following 8 hours of incubation in minimal media containing varying concentrations of cholesterol **d**, Western blot analysis of T389 S6K phosphorylation in Naïve B cell or B cells after 24 hours of stimulation with or without IL-4 in media with or without cholesterol, or 10% FBS +/-Rapamycin controls **e**, Mitochondrial mass measured using MitoTrackerTM Green of B cells following 72 hours of stimulation with or without IL-4 in media with or without cholesterol, or 10% FBS **f**, Representative gating and **g**, summary data of ER mass measured with ER-TrackerTM Green of B cells following 72H stimulation with or without IL-4 in minimal media supplemented M*β*CD-solubilized cholesterol, or 10% FBS and minimal media controls. Mean and S.D. representing 3 biological replicates on summarized plots. *P = 0.05, **P = 0.01, ***P < 0.001, ****P < 0.0001, unpaired two-tailed Student’s t-test. All experiments were performed at least twice.

Cholesterol canonically functions as a component of lipid rafts at the plasma membrane to support signaling, leading us to hypothesize that BCR signaling may be regulated by cholesterol availability. However, increasing anti-IgM concentrations to extreme hyperphysiological levels was insufficient to rescue proliferation *in vitro* in the absence of either IL-4 or cholesterol (Extended Data Fig.2a,b). To directly measure how cholesterol availability influenced BCR signaling strength, we incubated naïve B cells for 8 hours *ex vivo* in varying concentrations of cholesterol and subsequently labeled with INDO-1AM to flow cytometrically measure calcium flux in response to BCR ligation with anti-IgM Fab2 (Fig.3c). In line with recently published evidence that the pool of cholesterol in lipid rafts is metabolically inactive^22^, we observed no differences in BCR-induced calcium flux when exogenous cholesterol was limited.

Next, we hypothesized that cholesterol availability may influence the mTORC1 pathway. Phosphorylation of S6 kinase by mTORC1 occurs downstream of BCR/CD40, and stability of the mTORC1 complex has been shown to require lysosomal cholesterol in cell lines^23,24^. However, we did not observe any differences in S6K T389 phosphorylation with varied cholesterol availability after 24 hours of *in vitro* stimulation, further indicating that early signaling events are intact in these cells (Fig.3d). Consistent with this, cholesterol availability did not impair increases in cellular mitochondrial content following activation (Fig.3e), a process dependent on mTORC1 signaling^24,25^. Together, these data indicate IL-4/cholesterol signaling regulates G0/G1 progression and overall proliferative capacity of B cells via a mechanism that is downstream of BCR and mTORC1 signaling.

Since proximal signaling via IgM/CD40 was insensitive to cholesterol availability, we hypothesized that cholesterol may function as a downstream signaling molecule to license proliferation. The majority of the steps of cholesterol synthesis occur at the endoplasmic reticulum (ER), where the amount of cholesterol is highly regulated^12,26^. The ER is also the site where M*β*CD-solubilized cholesterol initially accumulates^27^, leading us to investigate how IL-4 and cholesterol impact ER expansion and functionality downstream of BCR signaling. By staining with ER-Tracker™ Green, a highly selective ER dye that binds sulphonylurea receptors of ATP-sensitive K+ channels prominent on ER, we found that accumulation of ER mass is sensitive to lipid deprivation and can be rescued by addition of either cholesterol or IL-4 (Fig.3f,g). This was not simply due to variation in total cell growth mitochondrial content was not affected by cholesterol or IL-4 (Fig.3e). Together, our data indicate that mitogenic signals require ER expansion to license G0/G1 progression and proliferation in B cells, that this regulatory step is sensitive to cholesterol synthesis at the ER, and that IL-4 signaling can alleviate reliance on exogenous cholesterol to promote B cell proliferation.

## Discussion

Transcriptional networks operate within biochemical parameters defined by the metabolic state of the cell; yet a clear understanding of how B cell fate is reciprocally regulated by gene expression and cell metabolism is lacking. We show here that mitogenic signals through BCR/CD40 require cholesterol in order to drive progression from G0 to G1 and initiate the proliferative phase of B cell activation. IL-4, a cytokine initially described as B cell growth factor, enables this process in the absence of extracellular lipids through lipid metabolic regulation at multiple scales. At the transcriptional level, IL-4 signaling activates STAT6 to upregulate expression of genes involved in cholesterol synthesis and downregulate expression of genes involved in cholesterol efflux to promote net accumulation of actively synthesized cholesterol. Together with SREBP2, this allows proliferation of cells in the absence of exogenous lipid nutrients, however how STAT6 engages SREBP2 at the molecular level to coordinately drive these transcriptional changes is unclear. Although the enzymes involved in cholesterol synthesis are upregulated, their activity is highly sensitive to substrate availability and product consumption by downstream enzymes. How this occurs downstream of IL-4 signaling will be important to explore in future work. Collectively, processes at multiple scales enable an accumulation of synthesized cholesterol in cells, driving expansion of the ER and licensing G0/G1 cell cycle progression.

Both G0/G1 progression as well as sensing organellar expansion are less understood than other modes of cell cycle regulation. Cell division requires doubling of cellular components in order to produce two functional daughter cells. Cell cycle checkpoints sensitive to DNA damage have been extremely well defined, however regulatory mechanisms sensitizing cells to defective or insufficient abundance of organelles are not as well understood. Recent work has defined an ER stress sensitive checkpoint in yeast cell cycle^28^. Our data suggest that, in mammalian cells, mitogenic signals require ER expansion to license G0/G1 progression and proliferation, and that this is sensitive to cholesterol content in the ER. We hypothesized that this process requires a threshold amount of cholesterol, and that IL-4 promotes endogenous cholesterol synthesis to relieve the reliance on exogenous cholesterol, which is likely limiting in microenvironments in secondary lymphoid tissues containing dense foci of proliferative cells. Here we put forward a mechanism through which cells evaluate lipid anabolic capacity via sensing actively synthesized cholesterol in the ER and, if metabolically equipped, expand the ER which enables progression through G0/G1 and entrance into cell cycle. Cholesterol is present at extremely low levels in the ER, rendering sensing mechanisms highly sensitive to small changes^26^. Our data are consistent with previous work connecting SCAP-dependent cholesterol synthesis with ER biogenesis and activation of CD8+ T cells^13,14^. How cholesterol is sensed in immune cells and what signaling components are engaged downstream remains an open area of investigation. Considering the critical known role that ER to nuclear signaling plays in B cell activation and differentiation, it will be important in future studies to define how the molecular dynamics of inter-organellar crosstalk license cell cycle entry and proliferation.

We demonstrate a physiological role for this axis in the context of early IL-4-dependent B cell response to influenza virus infection, where scRNA transcriptomics and trajectory analysis revealed a coincidence of IL-4 signaling and induction of programs to support net cholesterol accumulation with activation of B cells *in vivo*, which we show is also sensitive to statin treatment. This early expansion is necessary to provide a sufficient numbers of B cells for downstream diversification, selection, and differentiation, and imposes a bottleneck for the overall potential of the humoral immune response. A number of studies have suggested that statin treatment impairs humoral immune responses to Influenza vaccines in human patients^18–20^, however these results are controversial. Heterogeneity in human cohorts as well as confounding effects of long-term statin treatment on other cell types obscure any conclusions drawn about the direct effect of these drugs on B cells. Our work isolates a B cell intrinsic role for statin immune inhibition and defines one mechanism through which, at the cellular level, these drugs may impact vaccine strategies that by and large depend upon humoral immunity.

## Methods

### Mice

Mice were housed at a temperature of 72 °F (22°C) and humidity of 30–70% in a 12-h light/dark cycle with ad libitum access to food and water. Female mice aged 8–10 weeks at the start of the experiment were used throughout. Mice were bred and housed in the facilities of the Yale Animal Resources Center. C57BL/6N animals were purchased originally from Charles River and maintained in a breeding colony. C57BL/6J, B6.129S2(C)-Stat6tm1Gru/J (Stat6 Knockout), B6.SJL-Ptprca Pepcb/BoyJ (CD45.1), B6.C(Cg)-Cd79atm1(cre)Reth/EhobJ (Mb1-Cre), and Srebf2tm1Jdh/J (SREBP2 flox) were purchased from Jackson Laboratories.

### Infections

Mice were anesthetized i.p. with Ketamine/Xylazine (100 mg/kg body) and intranasally infected with 200 PFU of Influenza virus A/Puerto Rico/8/1934 (PR8) H1N1 strain expressing the LCMV gp33-41 epitope (PR8-33).1 Influenza virus was kindly provided by Drs. Susan Kaech and Jun Siong Low.

### B cell isolation

Single-cell suspensions were prepared from the spleens and lymph nodes of donor mice. Mature B2 B cells were enriched using immunomagnetic negative selection (StemCell Technologies) following a pre-incubation with biotin conjugated anti-CD9 (Biolegend Clone MZ3) and anti-CD93 (ThermoFisher Clone AA4.1) to deplete marginal zone and transitional B cells, respectively, at 5 *µ*g /mL for 20 minutes on ice. For proliferation analysis, isolated B cells were labeled at a concentration of 20 x 106 cells/mL in 1XPBS containing 5 *µ*M CellTraceTM Violet (Invitrogen C34557) for 12-14 minutes at 37C. The labeling reaction was quenched by washing with RPMI + 10% FBS and cells were washed twice with RPMI + 2 mM L-glutamine, 100 U/ml Pen Strep, and 55 *µ*M 2-Me prior to downstream culture.

### Cell culture

Isolated B cells were activated at a concentration of 2E6 cells/mL with 10 ng/mL rBAFF (R&D systems 8876-BF-01) and specified combinations of 5 *µ*g/mL (unless otherwise stated) goat anti-mouse IgM Fab2 (Jackson Immunoresearch 115-006-075), 1 *µ*g/mL anti-CD40 (bioxcell BP0016-2), and recombinant mouse Il-4 (peprotech 214-14).

Cell culture media contained RPMI+LG (Gibco 11875-093) supplemented with 10% FBS (Sigma F8192), 10% Dialyzed FBS (Gibco A3382001), or a cocktail containing Insulin, Transferrin, Selenium, Ethanolamine (Gibco 51500056) at a 1-100 dilution, and endotoxin-free recombinant human Albumin (Sigma A9731) at a final concentration of 5 mg/mL . All cell culture media contained 2 mM L-glutamine, 100 U/ml Pen Strep and 55 µ*µ*M 2-Me.

BSA-conjugated 16:0 palmitate and 18:1 oleic acid were generated as previously described and added at a concentration of 25 uM. M*β*CD-cholesterol (Sigma C4951) was added at a concentration of 4*µ*g/mL cholesterol unless otherwise specified. Thioridazine (Sigma T9025) was added at a final concentration of 5uM. Atorvastatin (Sigma PHR1422) was solubilized in DMSO and added at indicated concentrations.

### Flow cytometry

For labeling surface markers, single cell suspensions harvested from cell culture or isolated from spleen or mediastinal lymph nodes were incubated with 10% Rat serum (STEMCELL Technologies), Fixable Viability Dye (ebioscience) and surface antibodies. Intracellular stains were performed using commercially available Fix and permeabilization solutions coupled to incubation with Ki-67 and 0.5 µg /mL 7-AAD (Biolegend) or Pmp70/Abcd3.

For metabolic FACS analysis, cells were incubated at a final concentration of 50 *µ*g/ml 2-NBDG (Invitrogen N13195), 100 nM ER-Tracker GreenTM (Invitrogen E34251), or 40 nM MitoTrackerTM Green FM (Invitrogen M7514) in RPMI + 1% FBS for 30 minutes at 37C .

Multi-colour cytometry was performed on the LSR II flow cytometer (BD biosciences) or CytoFLEX S (Beckman Coulter) and analyzed with FlowJo v10.6.2.

### Calcium flux

Purified B cells were incubated in Minimal Media with varying concentrations of M*β*CD-solubilized cholesterol (Sigma C4951) for 8 hours at 37C in 5% CO2. 1 million cells were suspended in 400 *µ*L RPMI + 1%FBS, 10 mM Probenecid (Sigma P8761), 2 *µ*L 1.275 *µ*g /*µ*L Indo-1 AM (Sigma 13261), a Fixable Viability Dye (ebioscience), and an anti-B220 fluorophore conjugated antibody and kept at 30C throughout analysis.

Baseline fluorescence was recorded, and cells were then stimulated at a final concentration of 20 µg /mL goat anti-mouse Ig (Jackson Immuno 115-006-062). Ionomycin was used as a positive control. Cells were analyzed on an LSR II flow cytometer (Bd bioscience) and intracellular calcium levels were calculated using the kinetics function on FlowJo v.9.9.6.

### Immunofluoresence

Purified B cells were cultured for 72 hours in specified conditions. Cells were washed, resuspended at 0.5 E6/mL, and distributed onto slides centrifugally with Cytospin. Cells were fixed with 3% PFA, permeabilized with 0.5% Triton X-100, and stained with 1.5 ug/mL anti-Pmp70 (Sigma P0497) per the manufacturer’s protocol. Cells were stained with a secondary goat anti-Rabbit Cy3 (Invitrogen A10520) and DAPI (Sigma, F6057).

Confocal imaging was performed with a Nikon-Ti microscope combined with UltraVox spinning disk (PerkinElmer) and data were analysed by using the Volocity software (PerkinElmer).

### Western blot

Purified B cells were cultured for 72 hours in specified conditions. Negative controls were treated with 100 nM of Rapamycin (Sigma) at the onset of cell culture. Annexin V positive dead or dying cells were depleted using immunomagnetic negative selection (STEMCELL technologies).

Samples were normalized by cell number and lysed directly in RIPA Buffer before 1:1 dilution with 2x Lameli loading buffer. Protein extracts were separated on gradient 4% to 12% Bis–Tris SDS-PAGE gel (Invitrogen) and then transferred to a nitrocellulose membrane (Amersham). After blocking for 1 hour in a TBS containing 0.1% Tween 20 (TBST) and 5% nonfat milk, the membrane was probed with anti-phospho-T389 S6K1 (Cell Signaling 9234S clone 108D2) and anti-b-actin (Cell Signaling 4970 clone 13E5) diluted into TBST with 5% milk overnight. Membranes were washed 4x with TBST, followed by a 30-minute room temperature incubation with secondary antibodies conjugated to horseradish peroxidase diluted in TBST plus 5% milk. Membranes were washed as before and then developed using Pierce Supersignal West Femto detection kit and imaged with a Bio-Rad ChemiDoc imager.

### Quantitative PCR

Purified B cells were cultured for 72 hours in specified conditions. Dead or dying cells were depleted using immunomagnetic negative selection (STEMCELL technologies). RNA was isolated using a phenol-chloroform extraction and cDNA was generated using Bio-Rad iScript cDNA synthesis kit. Pre-validated primers were all ordered through Sigma KiqStart. Bio-Rad iTag Universal Sybr Green Supermix and Bio-Rad CFX6 rtPCR detection system were used for quantitative analysis. Fold change related to the control group was calculated using 2ΔΔCP method with Actb as the reference gene.

### Bulk RNA sequencing

For bulk RNA sequencing, purified B cells were cultured for 72 hours in specified conditions. Dead or dying cells were depleted using immunomagnetic negative selection (STEMCELL technologies) and RNA was extracted using Trizol and Qiagen RNeasy Kit. Technical duplicates were submitted to the Yale Center for Genome Analysis for cDNA synthesis and RNA sequencing. Libraries with unique barcodes were pooled and sequenced on Illumina NovaSeq S1 to produce 100 bp paired-end reads following manufacturer’s protocol. Raw reads from RNA-sequencing were aligned to the mouse genome mm10 using Kallisto2. Subsequently, differential expression analysis between groups was performed with DESeq^9^.

### Single cell RNA sequencing

For scRNAseq, single cell suspensions from mesenteric lymph nodes of 3 pooled mice on day 4 of PR8-gp33 Influenza infection were sorted on a Sony SH800 cell sorter. Populations of CD3+, CD19+, and CD3-CD19-CD45+ were pooled in a 1:1:1 ratio. Cells from pooled and approximately 10,000 cells were loaded onto a Chromium Controller (10x Genomics). Single-cell RNA-seq libraries were prepared using the Chromium Single Cell 5 v2 Reagent Kit (10x Genomics) according to manufacturer’s protocol and libraries were sequenced on the Illumina NovaSeq.

For data preprocessing, mRNA library reads were aligned to the mouse genome mm10 and quantified using CellRanger count4. Data was subsequently analyzed using Seurat (version 4.0)5–8. Data was first filtered to exclude cells in the 5th and 95th percentiles of library size and number of expressed genes, as well as cells with more than 5% mitochondrial transcripts, and include only genes expressed in more than 3 cells. Data was then normalized in terms of log (CPM/100 + 1), where CPM stands for counts per million (meaning that the sum of all gene-levels is equal 1,000,000). After normalization, we used ALRA (Adaptively-Thresholded Low Rank Approximation)^29^ to impute the matrix and fill in the technical dropout values. Subsequently, principal component analysis (PCA) was used to reduce data dimensionality followed by t-Distributed Stochastic Neighbor Embedding (t-SNE), a non-linear dimensionality reduction method. Louvain graph-based clustering was then used to generate clusters that were overlaid on the t-SNE coordinates to discern cell subpopulations.

Two individual clusters comprised B cells and were extracted into a new data subset. Monocle 2 was then used to perform semi-supervised ordering to specifically learn a trajectory defined by cellular activation. In a separate analysis, cells were subsetted based on imputed expression of CD19 which yielded similar results when subject to the same downstream trajectory analysis (data not shown). The normalized matrix output of ALRA was “de-normalized” by calculating the probability of expression for each gene where probability P is equal to the normalized expression value for each gene divided by the sum of normalized expression values for all genes in an individual cell; imputed counts were generated by using a multinomial model to sample count data for each cell using five times the original library size for each cell, and semi-supervised ordering in Monocle 2 was run using this sampled count data.

In Monocle 2, Btg2 and Myc were selected as “bellwether” genes to identify naïve and activated cells, respectively. The top 1000 genes that co-varied in either direction were determined and used to order cells along a developmental trajectory between these two states and assign a pseudotime value to each cell using reverse graph embedding.

### U^13^C-glucose labeling of cells and Isotope Enrichment Analysis

B cells activated under specified conditions were labeled in media containing 50% U13C-glucose. Cells were harvested at 72 hours and Annexin V+ dead/dying cells were depleted using immunomagnetic negative selection (STEMCELL technologies). Isotopomeric Spectral Analysis (ISA) was performed as previously described^14^.

### Statistical analysis

Experiments were conducted with technical and biological replicates at an appropriate sample size as estimated by our prior experience. No statistical methods were used to predetermine sample size. No methods of randomization and no blinding were applied. All data were replicated independently at least once as indicated in the figure legends, and all attempts to reproduce experimental data were successful. For all bar graphs, mean and S.D. are shown. All statistical analysis was performed using GraphPad Prism 7 or newer. p-values <0.05 were considered significant (*p<0.05, **p<0.01; ***p<0.001, **** p<0.0001); p-values >0.05; non-significant (ns). All sample sizes and statistical tests used are detailed in figure legends.

## Acknowledgements

We would like to thank J. Alderman, C. Lieber, L. Evangelisti, C. Hughes, and E. Hughes-Picard for technical and administrative assistance. We would like to thank Dr. V. D. Dixit, Dr. S. Kaech, Dr. J. Pereira, and Dr. R. Jackson for helpful comments and discussion, and G. Linderman for advice on computational analysis. H.S. is supported by the National Science Foundation Graduate Research Fellowship Program under program grant number Fall 2007 DGE1752134. W.B. was supported by a K22 (NIAID 5K22AI141758. This work was supported by NIH grants R37AR40072 (J.C. and R.A.F) and R01AR074545 (J.C.), by the Lupus Research Alliance (J.C.), and by the Howard Hughes Medical Institute (R.A.F and A.Y.).

## Author contributions

Conceptualization and design: HRS, AGY Data acquisition and analysis-mass spectrometry: HRS, AGY, KJW, QZ, YH Data acquisition and analysis-immunofluorescence: HRS, JRB Data acquisition and analysis-scRNAseq: HRS, AGY, JZ, RQ, MHS, YK Data acquisition and analysis-qPCR: HRS, AGY Data acquisition and analysis-all else: HRS Technical support and suggestions: JZ, KJW, QZ, YH, JRB, RQ, MHS, JAS, CH, ES, WKM, WB, CC Writing-original draft: HRS Writing-review and edit: HRS, AGY, MHS, JAS, CH, WKM, CC, SJB, JC, RAF Project Administration: SJB, JC, RAF

## Competing interests

R.A.F. is a scientific advisor to GlaxoSmithKline. All other authors declare no competing interests.

**Supplementary Figure 1.**
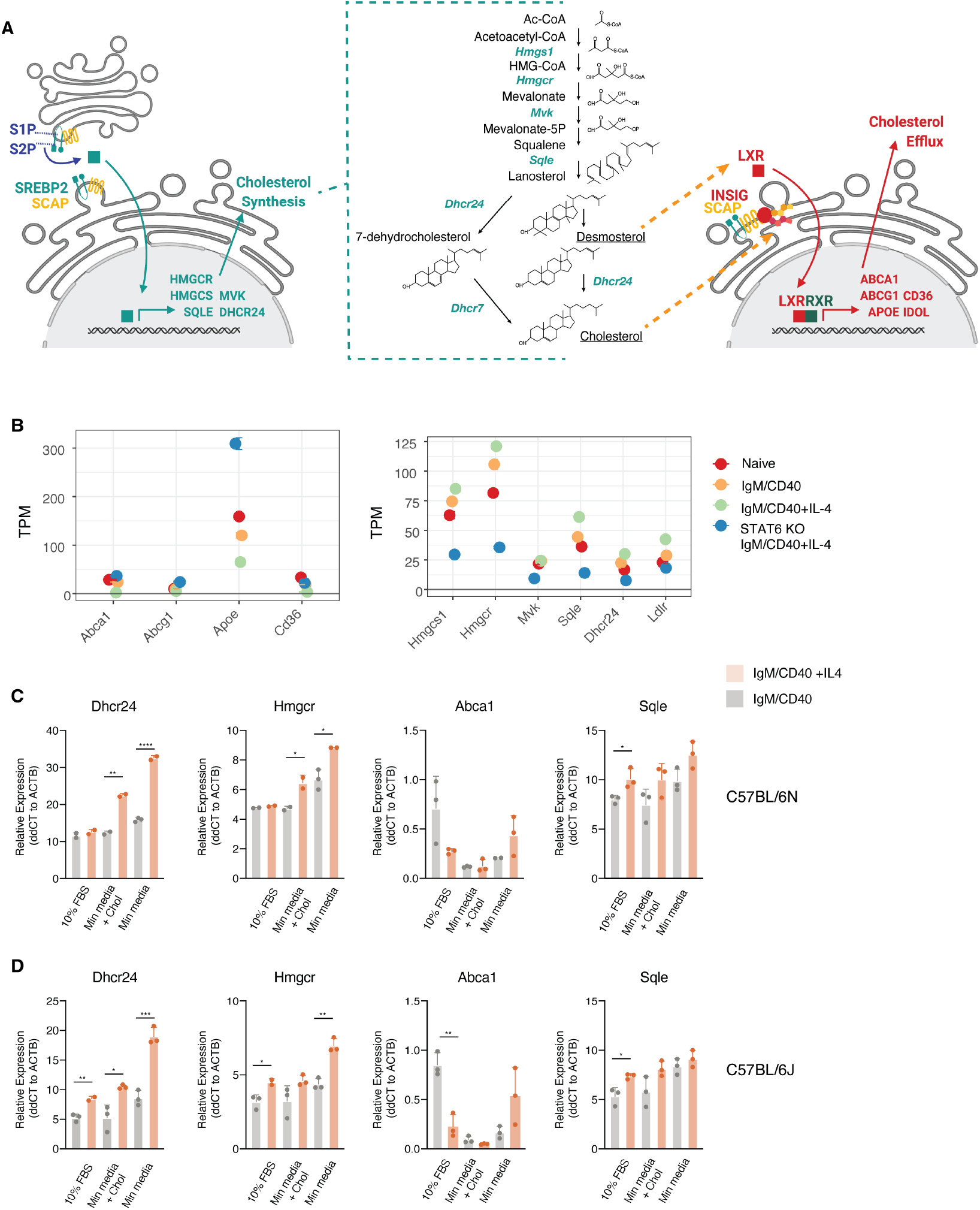
IL-4 regulates cholesterol synthesis through transcriptional reprogramming but does not affect glucose uptake. **a**, Graphical illustration of cholesterol homeostatic regulation by SREBP2 and LXRs **b**, Transcript abundance of genes involved in cholesterol synthesis, cholesterol efflux, and IL-4 signature in Naïve cells, cells activated for 72H with anti-IgM/CD40, and either wild type or STAT6 KO cells activated with anti-IgM/CD40 + IL-4 **c**, qPCR validation of gene expression changes in B cells from C57BL6/N and c, C57BL6/J in response to IL-4 treatment following 72H activation. Mean and S.D. from 3 technical replicates are represented on summarized plots. **P = 0.01, unpaired two-tailed Student’s t-test used to determine significance.

**Supplementary Figure 2.**
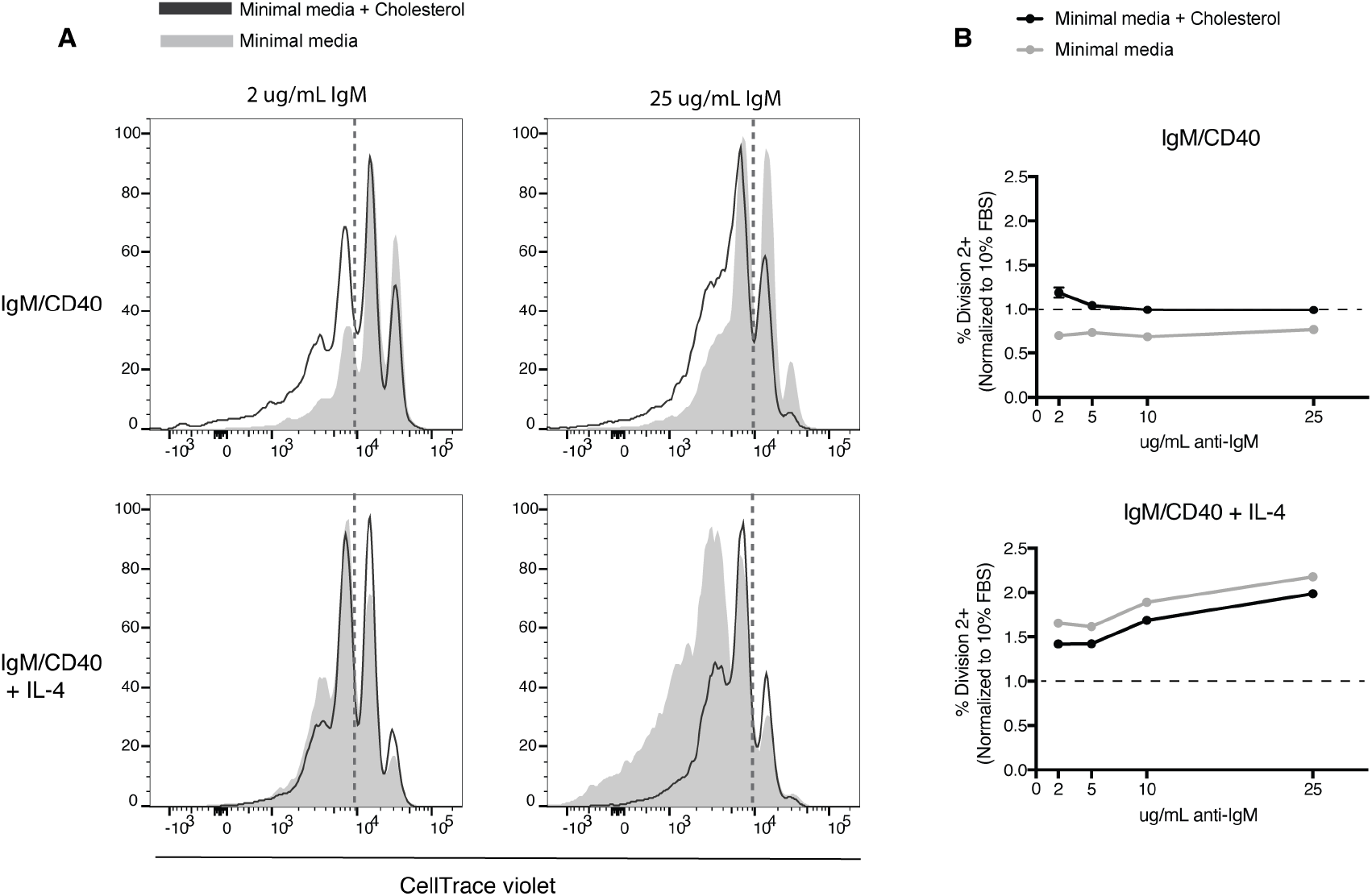
Intensified BCR signaling is insufficient to rescue proliferation in the absence of IL-4 or exogenous cholesterol. **a**, Proliferation of B cells activated for 96H with low (2 *µ*g/mL) or high (25 *µ*g/mL) amounts of anti-IgM with anti-CD40 +/-IL-4 **b**, Proliferation of B cells activated for 96H with varying amounts of anti-IgM with anti-CD40 +/-IL-4

